# Accumulation of Tau in Extracellular Vesicles Disturbs the Astrocytic Mitochondrial System

**DOI:** 10.1101/2023.02.15.527595

**Authors:** Romain Perbet, Valentin Zufferey, Elodie Leroux, Enea Parietti, Jeanne Espourteille, Lucas Culebras, Sylvain Perriot, Renaud Du Pasquier, Séverine Bégard, Vincent Deramecourt, Nicole Déglon, Nicolas Toni, Luc Buée, Morvane Colin, Kevin Richetin

## Abstract

Tauopathies are neurodegenerative disorders involving the accumulation of tau isoforms in cell subpopulations such as astrocytes. The origins of the 3R and 4R isoforms of tau that accumulate in astrocytes remain unclear.

Extracellular vesicles (EVs) were isolated from primary neurons overexpressing 1N3R or 1N4R tau or from human brain extracts (progressive supranuclear palsy or Pick disease patients or controls) and characterized (electron microscopy, nanoparticle tracking analysis (NTA), proteomics). After the isolated EVs were added to primary astrocytes or human iPSC-derived astrocytes, tau transfer and mitochondrial system function were evaluated (ELISA, immunofluorescence, MitoTracker staining). We demonstrated that neurons in which 3R or 4R tau accumulated had the capacity to transfer tau to astrocytes and that EVs were essential for the propagation of both isoforms of tau. Treatment with tau-containing EVs disrupted the astrocytic mitochondrial system, altering mitochondrial morphology, dynamics and redox state. Although similar levels of 3R and 4R tau were transferred, 3R tau-containing EVs were significantly more damaging to astrocytes than 4R tau-containing EVs. Moreover, EVs isolated from the brain fluid of patients with different tauopathies affected mitochondrial function in astrocytes derived from human iPSCs. Our data highlight that tau pathology spreads to surrounding astrocytes via EVs-mediated transfer and modify their function.

## 1. Introduction

Tauopathies are a group of more than 20 diseases that include Alzheimer’s disease (AD), progressive supranuclear palsy (PSP), Pick disease (PiD), frontotemporal lobar degeneration and primary age-related tauopathy.

Although all tauopathies are associated with the accumulation of tau (predominantly in neurons), there are notable differences among them in (1) the affected region, (2) the presence or absence of tau inclusions in glia and (3) the tau isoform (3R/4R) that constitutes the observed inclusions/aggregates. Indeed, as a result of alternative splicing, six major isoforms of tau, each with a microtubule-binding region consisting of 3 (3R tau isoforms) or 4 (4R tau isoforms) repeated sequences, coexist in the human brain [1,2]. In the healthy human brain, the 3R and 4R tau isoforms are present at equally low levels in glial cells [3-5]. However, analysis of patients’ brains revealed that some tauopathies are exclusively associated with 3R or 4R tau inclusions, whereas some are associated with mixed tau inclusions [6]. Additionally, many studies have shown the presence of tau inclusions in glial cells, including astrocytes, oligodendrocytes and microglia [7-10].

Also, there is evidence of prion-like propagation of tau pathology, especially in AD [11]. This process involves neurons and affects glial cells [12]. While the origin of tau in these cells is not well defined, astrocytes are a particularly interesting possibility due to their function in the tripartite synapse [13]. Tau uptake and its role in tau pathology spreading through astrocytes are now the subject of increasing attention [14]. De Gérando and collaborators showed that astrocytic tau pathology can emerge secondary to neuronal pathology [15]. It also appears that astrocytes can internalize tau fibrils, which are subsequently degraded by lysosomes, potentially contributing to reduced tau spreading [16]. The nature of these tau species and how they are shuttled from neurons to astrocytes are currently unknown.

Considering the importance of 1) tau propagation in interconnected regions and 2) the role of astrocytes in synaptic function, it is reasonable to study the role of astrocytes in tau propagation and the underlying mechanisms. A few studies have investigated tau transfer between neurons and astrocytes in cell and animal models [15,16], but the cellular mechanisms underlying this exchange of material are unexplored, and no data from human samples are available. Cells exchange material with their neighbors in various ways, and these intercellular communications are fundamental to maintaining tissue homeostasis, which is often dysregulated in disorders. Extracellular vesicles (EVs) have emerged as critical cell-to-cell communication regulators under physiological conditions and in diseases such as neurodegenerative diseases [17,18]. They are secreted by cells through unconventional protein secretion [19] and shuttle many components, such as nucleic acids, lipids, active metabolites, and cytosolic and cell surface proteins [20]. Due to their membrane composition, they should act as unique intercellular delivery vehicles for the transfer of pathological species between cells to allow the propagation of tau pathology.

We and others recently demonstrated that EVs isolated from AD brain-derived fluids (BDFs) play a role in human tau spreading [21,22]. While tau has been detected in exosomes isolated from the primary astrocyte cultures [23], their involvement in tau propagation is still debated. Others have suggested that tau is mainly found in exosomes derived from microglia from animal models [9]. Whether tau accumulation in astrocytes is beneficial or deleterious to astrocytes remains also to be elucidated. We recently observed complex topographic patterns of tau isoforms and accumulation of amyloid-β in the hippocampi of AD patients. We show that the accumulation of 3R tau (but not phospho-4R tau) in astrocytes is exacerbated by amyloid-β accumulation and impairs mitochondrial function and ATP production, thus inducing a reduction in the number of inhibitory neurons in the hippocampus, which is associated with memory decline [24]. However, the mechanism of 3R and 4R tau accumulation and their origin in hippocampal astrocytes are still poorly understood. We wondered whether EV-mediated transfer of 3R/4R tau might underlie tau accumulation in astrocytes and the consequent astrocyte dysfunction. In the present work, we demonstrated that **1)** 3R and 4R tau are transferred from neurons to astrocytes, **2)** while neuronal tau is mainly secreted in the free form, tau isoforms are shuttled from neurons to astrocytes mainly through EVs (large EVs), and **3)** EVs-mediated tau accumulation, especially 3R tau accumulation, in astrocytes alters mitochondrial function.

## 2. Materials and methods

### 2.1. Human samples

Prefrontal brain extracts from non-demented control subjects (n = 5), PSP (n = 5) and PiD (n = 5) patients were obtained from the Lille Neurobank (fulfilling French legal requirements concerning biological resources and declared to the competent authority under the number DC-2008-642); donor consent and ethics committee approval were obtained, and the data were protected. The samples were managed by the CRB/CIC1403 Biobank, BB-0033-00030. The demographic data are listed in Table 1.

**Table 1.**
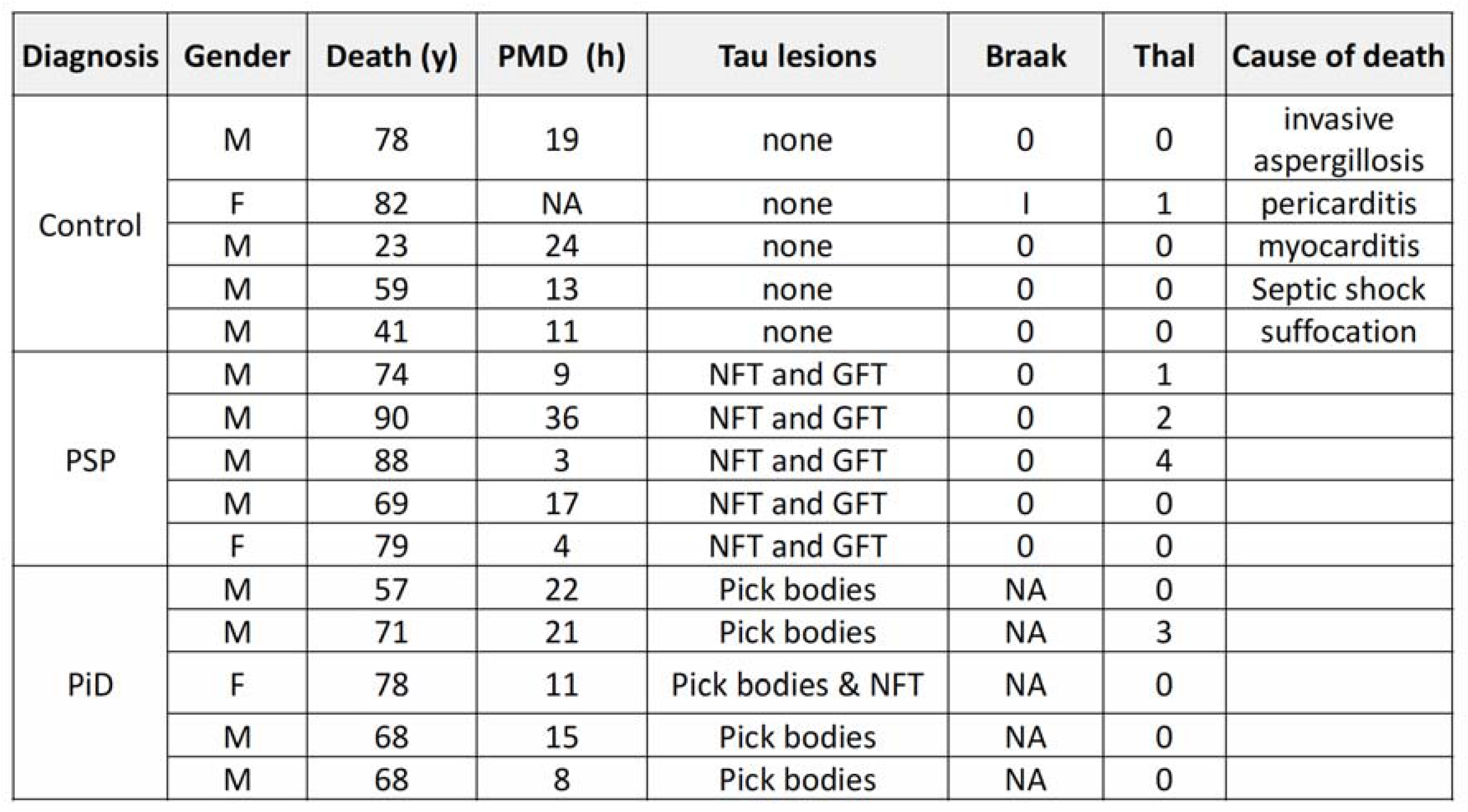
Demographic, biological, and clinical characteristics of the brain sample donors. The characteristics of the donors who provided brain samples used for BDF isolation are listed (n = 5 non-demented control subjects, n = 5 PSP patients and n = 5 PiD patients). PMD, postmortem delay; NFTs, neurofibrillary tangles, GFTs, glial fibrillary tangles; NA, not applicable.

| Diagnosis | Gender | Death (y) | PMD (h) | Tau lesions | Braak | Thal | Cause of death |
| --- | --- | --- | --- | --- | --- | --- | --- |
| Control | M | 78 | 19 | none | 0 | 0 | invasive aspergillosis |
|  | F | 82 | NA | none | 1 | 1 | pericarditis |
|  | M | 23 | 24 | none | 0 | 0 | myocarditis |
|  | M | 59 | 13 | none | 0 | 0 | Septic shock |
|  | M | 41 | 11 | none | 0 | 0 | suffocation |
| PSP | M | 74 | 9 | NFT and GFT | 0 | 1 |  |
|  | M | 90 | 36 | NFT and GFT | 0 | 2 |  |
|  | M | 88 | 3 | NFT and GFT | 0 | 4 |  |
|  | M | 69 | 17 | NFT and GFT | 0 | 0 |  |
|  | F | 79 | 4 | NFT and GFT | 0 | 0 |  |
| PiD | M | 57 | 22 | Pick bodies | NA | 0 |  |
|  | M | 71 | 21 | Pick bodies | NA | 3 |  |
|  | F | 78 | 11 | Pick bodies & NFT | NA | 0 |  |
|  | M | 68 | 15 | Pick bodies | NA | 0 |  |
|  | M | 68 | 8 | Pick bodies | NA | 0 |  |

### 2.2. Rat primary neuron and astrocyte cultures

Primary hippocampal neurons were prepared from 17-days-old Wistar rat embryos, as previously described [25]. Five days later, the cells were infected with a lentiviral vector (LV) encoding human 1N4R-V5 or 1N3R-V5 (PGK-4R-V5-SIN, PGK-3R-V5-SIN), as previously described [26]. The LVs were produced and validated as previously described [27,28].

Primary hippocampal astrocytes were isolated from postnatal day 1 rat pups (Wistar rats from Janvier Laboratory) as previously described [29]. Briefly, the brains were removed aseptically from the skulls, the meninges were excised under a dissecting microscope, and the hippocampus was dissected. The cells were dissociated by passage through needles of decreasing gauge (1.1 × 40; 0.8 × 40 and 0.5 × 16) 4 or 5 times with a 5-ml syringe. The cells were plated at a density of 20,000 cells per cm^2^ in 6-well plates in DMEM containing 25 mM glucose and supplemented with 10% fetal calf serum, 44 mM NaHCO_3_, and 10 ml/L antibiotic/antimycotic solution (pH 7.2) in a final volume of 3 ml/well and incubated at 37°C in an atmosphere containing 5% CO_2_/95% air. The culture medium was replaced 4–5 days after plating and every 2-3 days thereafter after gently tapping the plates to remove the less adherent cells (oligodendrocytes and microglia). For the neuron-to-astrocyte tau transfer experiment, 10×10^5^ astrocytes were treated with 10 µl of neuron-derived small EVs (ND-SEVs), neuron-derived large EV (ND-LEVs) and neuron-derived free protein (ND-FFP) (obtained from 10×10^6^ neurons) at days *in vitro* 10 (D.I.V. 10). For the microfluidic experiment, astrocytes plated in 6-well plates were detached with trypsin at D.I.V. 10 and plated in the axonal compartment. To monitor the effects of tau accumulation in astrocytes on astrocytic mitochondria, cells were infected with a LV encoding Mitotimer (LV-G1-MitoTimer-miR124T) at D.I.V. 7-8, as previously described [26].

### 2.3. Human iPSC-derived astrocyte culture

Human iPSCs were differentiated into astrocytes as previously described in detail [30]. Briefly, human iPSCs were plated onto poly-L-ornithine/laminin (PO/L)-coated plates and cultured in neural induction medium (DMEM/F-12 supplemented with N2 supplement (1x), B27 supplement without vitamin A (1x), noggin (500 ng/ml), SB431542 (20 μM) and FGF2 (4 ng/ml)) to allow the formation of embryoid bodies. The medium was changed every other day for 12 days. After 12 days, neural precursor cells (NPCs) were passaged and grown for at least 3 weeks in single-cell culture in NPC expansion medium (DMEM/F-12 supplemented with N2 supplement (1x), B27 supplement without vitamin A (1x), FGF2 (10 ng/ml) and EGF (10 ng/ml)). Next, the medium was replaced with astrocyte induction medium (DMEM/F-12 supplemented with N2 supplement (1x), B27 supplement without vitamin A (1x), LIF (10 ng/ml) and EGF (10 ng/ml)). The medium was changed every other day, and the cells were passaged when they reached confluence. After 2 weeks, the medium was replaced with astrocyte medium (DMEM/F-12 supplemented with B27 supplement without vitamin A) supplemented with CNTF (20 ng/ml). After culture for 4 weeks in this medium, the cells displayed the cellular characteristics of mature human astrocytes [31]. Then, the astrocytes were infected with a LV encoding Mitotimer (LV-G1-MitoTimer-miR124T). Four days later, the astrocytes were treated with 10 µl of BDF-LEVs (10^6^ EVs per astrocyte), and mitochondrial system function and dynamics were evaluated. The dose of EVs was selected based on a cell toxicity experiment (the number of DAPI-stained cells remaining 24 h after treatment was counted and normalized to that in the control condition); in this experiment, no toxicity was observed at this dose (**Figure S1**). These experiments were performed according to a protocol approved by our institutional review committee (approval n° CER-VD 2018-01622).

### 2.4. Microfluidic system and immunofluorescence

Glass coverslips were coated overnight at 4°C with 0.5 mg/ml poly-D-lysine. Microfluidic chambers (AXIS™, Temecula, CA) were subsequently placed on the coated glass coverslips and attached to the glass. All chambers had microgrooves 450 μm in length and 10 μm in width. Rat primary embryonic neurons were cultured as described above, and approximately 30,000 cells were plated in the two wells of the somatodendritic compartment. The cultures were maintained at 37°C for 7 days to allow differentiation, with a volume gradient from the somatodendritic compartment to the astrocytic compartment to help axonal guidance. At D.I.V. 7, Tau-V5-LV (200 ng of LV per well) was added to the somatodendritic compartment after first reversing the volume gradient between the compartments to counteract viral diffusion. The quality controls for compartment isolation were previously described by Dujardin and collaborators [26]. Primary astrocytes were then plated in the axonal compartment in neurobasal medium and maintained at 37°C for 5 days. V5 immunolabeling was performed to evaluate tau transfer from neurons to astrocytes. The cells in the compartments (somatodendritic and axonal) were washed once with phosphate-buffered saline (PBS) and fixed with 4% paraformaldehyde (PFA) for 20 min. After removing the fixative, the cells were washed three more times with 50 mM NH4Cl and permeabilized with Triton X-100 (0.1%, 10 min at room temperature). Subsequently, the slides were incubated with primary antibodies at 4°C overnight, and labeling was performed by incubation with the appropriate Alexa Fluor-conjugated secondary antibodies (1:400) for 45 min at room temperature. The cells were mounted with Vectashield medium containing DAPI.

### 2.5. EVs isolation from primary cultures

Cells (10×10^6^) in fresh medium were collected and placed on ice 10 days after LV infection. Protease inhibitors were added before centrifugation for 10 min at 2,000 × g and 4°C. The supernatant was centrifuged for 50 min at 20,000 × g, and the pellet, which contained ND-LEVs, was collected. The supernatant was centrifuged for 50 min at 100,000 × g to obtain ND-SEVs (pellet) and FFP (supernatant). ND-LEVs and ND-SEVs were suspended in 100 µl of RIPA buffer (150 mM NaCl, 1% NP40, 0.5% sodium deoxycholate, 0.1% SDS, and 50 mM Tris HCl; pH= 8.0) for biochemical assays, 100 µl of 4% PFA (diluted in phosphate buffer (0.08 M Na2HPO4 and 0.02 M NaH2PO4)) for electron microscopy analyses or 100 µl of phosphate buffer (0.08 M Na2HPO4 and 0.02 M NaH2PO4) for astrocyte treatment. The FFP was concentrated to a volume of 100 µl with an Amicon device (3 kDa).

### 2.6. EVs isolation from human prefrontal cortex tissue

BDF-EVs were isolated from human prefrontal cortex tissue as previously described [32]. Briefly, the tissue was incubated on ice in 5 ml of Hibernate-A. It was gently mixed in a Potter homogenizer, and 2 ml of 20 units/ml papain in Hibernate-A was added to the homogenate for 20 min at 37°C with agitation. Then, 15 ml of cold Hibernate-A buffer (50 mM NaF, 200 nM Na_3_VO_4_, 10 nM protease inhibitor) was added, and the sample was mixed by pipetting to stop the enzymatic activity. Successive centrifugation was performed at 4°C (300, 2000, and 10 000 × g) to remove cells, membranes, and debris, respectively. The final supernatant was kept at -80°C until EVs isolation was performed. BDF-LEVs and BDF-SEVs were pooled from 5 patients per group. EVs isolation from cell medium or human BDF was carried out using differential ultracentrifugations as previously described [33] to obtain LEVs-, SEVs- and FFP-enriched fractions. The final pellets were suspended in a final volume of 400 µl and kept at -80°C.

### 2.7. Nanoparticle tracking analysis (NTA)

The size, number and distribution of particles were determined using NTA (Nanosight NS300, Malvern) as described previously [22]. To generate statistical data, five 90-second videos were recorded and analyzed using NTA software (camera level: 15; detection threshold: 4).

### 2.8. Electron microscopy [22,33]

Samples (5 μl) were deposited on a carbon film-supported grid (400 mesh) and incubated at room temperature (RT) for 20 min. The grids were fixed in PBS-glutaraldehyde (1%) for 5 min at RT and then rinsed in distilled water. They were incubated for 5 min in 1% uranyl acetate and for 10 min on ice in a mixture of 1% uranyl acetate/2% methylcellulose. Dry grids were observed under a transmission electron microscope (Zeiss EM900). When indicated, immunolabeling was performed. The grids were rinsed once in PBS and incubated twice (3 min at RT) in PBS-50 mM glycine before incubation in PBS-1% bovine serum albumin (BSA) for 10 min at RT. A primary antibody diluted in PBS-1% BSA was applied for 1 h at RT and detected using an appropriate secondary antibody diluted in PBS-1% BSA (18 nm gold colloidal goat anti-mouse). After rinsing in PBS, the grids were processed as described above.

### 2.9. Antibodies

The following antibodies were used for immunohistofluorescence and immunohistochemistry at the indicated dilutions: a polyclonal rabbit antibody against the C-terminus of Tau (C-Ter), which recognizes the last 15 AA of the protein (C-Ter, raised in-house, 1:1000 for electronic microscopy) [34], A mouse monoclonal antibody against V5 (1:10000) that recognizes the V5 epitope of tagged tau [24,26], and a polyclonal rabbit antibody that recognizes glial fibrillary acidic protein (GFAP) (1:10000).

### 2.10. ELISA

Fractions were obtained after ultracentrifugation of culture medium or BDF as described above. Tau levels were determined using INNOTEST hTau Ag (Fujirebio/Innogenetics, Belgium), which is a sandwich ELISA microplate assay for the quantitative determination of human tau antigen levels in fluids, according to the manufacturer’s instructions. The capture antibody was an AT120 antibody, and biotinylated HT7 and BT2 antibodies were used as detection antibodies [35]. For cell samples, we customized the INNOTEST hTau Ag ELISA kit to quantify Tau-V5 levels. The original INNOTEST kit plate was replaced with a 96-well MicroWell™ MaxiSorp™ flat bottom plate coated with V5 antibody (Invitrogen P/N-0705; 1:10000) in carbonate buffer (NaHCO_3_ 0.1 mM, Na2CO_3_ 0.1 mM; pH=9.6) overnight at 4°C). The other steps were performed according to the manufacturer’s instructions. Tau uptake by astrocytes was evaluated 24 h after incubation of astrocytes with ND-FFP and ND-LEVs fractions from control neurons (NI) or neurons overexpressing 1N3R or 1N4R Tau-V5. Tau uptake was calculated with the following equation: ([Tau-V5] secreted by neurons / [Tau-V5] in astrocytes) x 100.

### 2.11. Quantification of Tau-V5 levels in primary culture

The optical density (OD) of Tau-V5 was measured in 5 regions of interest (ROI) in each compartment (somatodendritic compartment, microgrooves, and axonal compartment). The optical density in each region of interest was evaluated using Zen 2 image analysis software (blue edition). Each OD value was normalized by subtracting the OD in an ROI in which no Tau-V5 signal was present (NI). The OD of Tau-V5 in astrocytes was measured from confocal scanning images taken with a Zeiss LSM 710 Quasar microscope equipped with a 40× oil immersion objective at a z-axis resolution of 0.9 µm. We used Imaris software to measure the intensity of Tau-V5 in the reconstructed GFAP+ region of the axonal compartment.

### 2.12. Analysis of mitochondrial dynamics by Mitotimer

The complete method was described in previous publications [36,37]. Briefly, using an inverted microscope (Nikon Eclipse Ti-2) (150x magnification, 100x oil immersion objective, 1.5x intermediate magnification) and the Perfect Focus System (PFS), 16 bit image sequences (1 frame/s for 60 s) were taken at baseline (BL) and 6 h and 24 h after treatment with EVs. Sequential excitation at 490 nm (for the green channel) and 550 nm (for the red channel) and detection of green (500-540 nm) and red (550-600 nm) signals were performed. Then, using GA3 software, we selected the first frame of each image sequence in the red and green channels to generate binary masks for each mitochondrion. Then, mitochondrial features (red/green ratio, elongation, area, length, branches, and junctions) were automatically measured for each mitochondrion and averaged for each astrocyte. We performed these analyses for at least 20-25 cells per condition (a minimum of fifty mitochondria per cell). Manual motility analysis is preferred for dynamic and mobility criteria (displacement, track length, speed, straightness, and events) due to the high complexity of mitochondrial movements. Nikon’s NIS Element system was used to manually track 25-50 mitochondria per cell. Finally, after log transformation, the change in score [37] relative to the BL was calculated and normalized to that of the control group (ND-LEVs-CFP for primary cultures and BDF-LEVs from human controls) [36].

### 2.13. Sample sizes, calculations and statistical analysis

Human samples were classified based on neurological and neuropathological examination. The order of culture and procedures was randomized for each experiment. Investigators were blinded to group allocation when processing the tissue and performing cell counts and during confocal image acquisition. Values are presented as the mean□±□s.e.m.; N corresponds to the number of independent experiments, and n corresponds to the overall number of values. Statistical analyses of raw data were performed with GraphPad Prism software v8.0. The normality of the data was verified using the Shapiro□Wilk test. Differences between two experimental groups were analyzed using Student’s t test (parametric test) or the Mann–Whitney test (nonparametric test). Statistical analyses of differences among more than two experimental groups were performed using one-way ANOVA followed by Dunn’s post hoc analyses for multiple comparisons (parametric test) or the KruskalLWallis test (nonparametric test). For Mitotimer score change analysis, as previously described [36], we used two-way ANOVA followed by Tukey’s post hoc analyses to compare the differences in mitochondrial function between the 3R and 4R tau groups with time as the independent variable. In addition, to compare each parameter measured at 6 h and 24 h to the baseline values for each astrocyte, we used the Wilcoxon signed-rank test.

## 3. Results

### 3.1. 3R and 4R tau transfer from neurons to astrocytes

To study the origin of the accumulated 3R and 4R tau in astrocytes, we overexpressed human wild-type 1N3R and 1N4R tau (fused to a V5 tag) in primary hippocampal neuron cultures in a microfluidic system. Due to axonal growth in the microgrooves, the hippocampal neurons in the somatodendritic compartment (SD) were connected with a second chamber (axonal compartment, AX) containing primary astrocytes. 1N3R or 1N4R tau was overexpressed in the somatodendritic compartment using LVs (Figure 1A). Five days later (D.I.V. 12), immunochemistry for V5 revealed accumulation of 1N3R and 1N4R tau in the somatodendritic (containing soma and dendrites) compartment and axons in the microgrooves (Figure 1B). Both isoforms were overexpressed in neurons at similar levels (Figure 1C). Interestingly, we observed that a large population of astrocytes in the axonal compartment (identified by GFAP immunochemistry) exhibited Tau-V5 puncta in the cytoplasm (Figure 1B, D). Quantification of the number of 1N3R and 1N4R tau inclusions in astrocytes through measurement of the V5 intensity in GFAP+ astrocytes indicated that both isoforms could be transferred from hippocampal neurons to astrocytes to similar degrees (Figure 1E).

**Figure 1.**
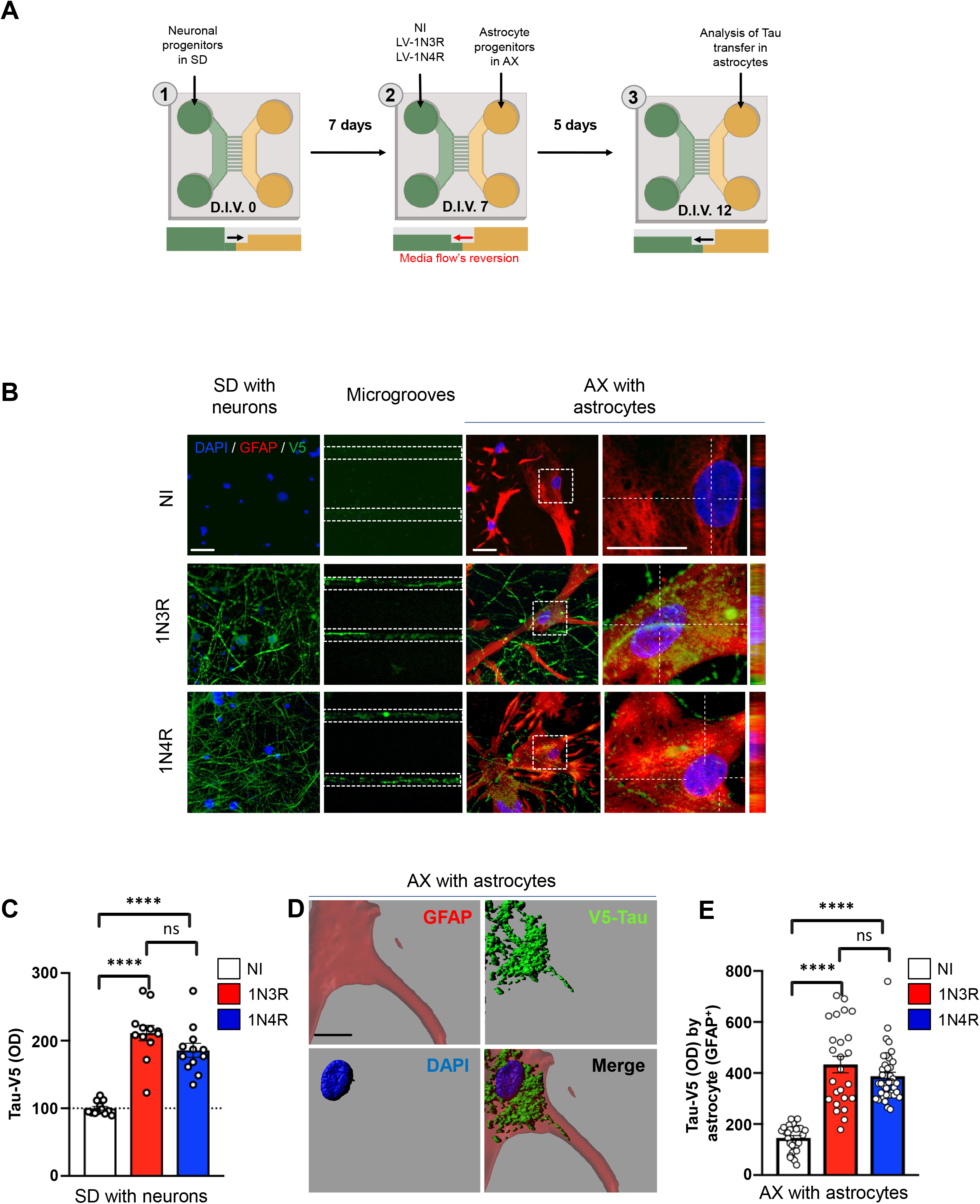
Tau is shuttled from neurons to astrocytes. **(A)** Schematic of the microfluidic system used to investigate tau transfer from hippocampal neurons to astrocytes. **(B)** Confocal micrographs showing 1N3R or 1N4R tau (V5+) and astrocytes (GFAP+) in the somatodendritic and axonal compartments and microgrooves. **(C)** Histogram showing the Tau-V5 optical density in the somatodendritic compartment. **(D)** 3D reconstruction of confocal micrographs showing tau (V5+) in an astrocyte (GFAP+) in the axonal compartment. **(E)** Histogram showing the number of Tau-V5 inclusions detected per astrocyte. The scale bars are 20□µm **(B)** and 5 µm **(D)**. SD= somatodendritic compartment, AX= axonal compartment, LV=lentiviral vector, NI= noninfected. For **C** and **E**, N□=□cultures/microfluidic chambers/cells: NI: 1/3/32, 1N3R: 1/3/25, 1N4R :4/3/39. Ordinary one-way ANOVA with Sidak’s multiple comparison test.

### 3.2. Neuronal tau is mainly secreted in the free form

Transfer of tau to adjacent cells requires secretion by the donor cell. First, we assessed which form(s) of tau is (are) secreted by primary neurons. Culture medium from primary neurons overexpressing 1N3R or 1N4R tau at similar levels (Figure 2A, B) was collected and fractioned to separate EVs from free proteins. EVs present in extracellular media are a heterogeneous population in which different biogenesis pathways are active. EVs include (1) exosomes (small vesicles, size <150 nm), which are generated from multivesicular bodies containing intraluminal vesicles that are secreted into the extracellular fluid, and (2) ectosomes (large vesicles, size>150 nm) that originate from direct plasma membrane budding [20]. To further investigate the EV subtypes that might be involved in tau transfer to astrocytes, we separated small from large vesicles as previously reported [33]. Fractions containing ND-LEVs, ND-SEVs and ND-FFP were then collected and characterized. Electron microscopy (Figure 2C) and NTA (Figure 2D, E) confirmed that ND-SEVs and ND-LEVs with intact structures were present in our fractions. A total of 26.3 ± 5.8% and 35 ± 18.7% of ND-SEVs with a size > 150 nm and 73.6 ± 5.8% and 65 ± 18.7% of ND-SEVs with a size between 10 and 150 nm have been isolated from cells overexpressing 1N3R and 1N4R tau, respectively (Figure 2D). A total of 71.3 ± 9.2% and 63.6 ± 13.8% of ND-LEVs with a size > 150 nm and 28.6 ± 9.2% and 36.3 ± 13.8% of ND-LEVs with a size between 10 and 150 nm have been isolated from cells overexpressing 1N3R and 1N4R tau, respectively (Figure 2E). It should be noted that size repartition was not altered by the expression of different tau isoforms in neurons (Figure 2D, E nonsignificant difference between the 1N3R and 1N4R groups). The presence of tau in these fractions was detected by electron microscopy (Figure 2F), and tau levels were quantified by ELISA (Figure 2G). While a large majority (93% ± 0.2 and 93.3 ± 0.2 for 1N3R and 1N4R tau, respectively) of tau protein was found in the free form (ND-FFP), only 4.7% ± 0.3 of 1N3R tau and 4.5% ± 0.2 of 1N4R tau was found in the ND-LEVs fractions (Figure 2G).

**Figure 2.**
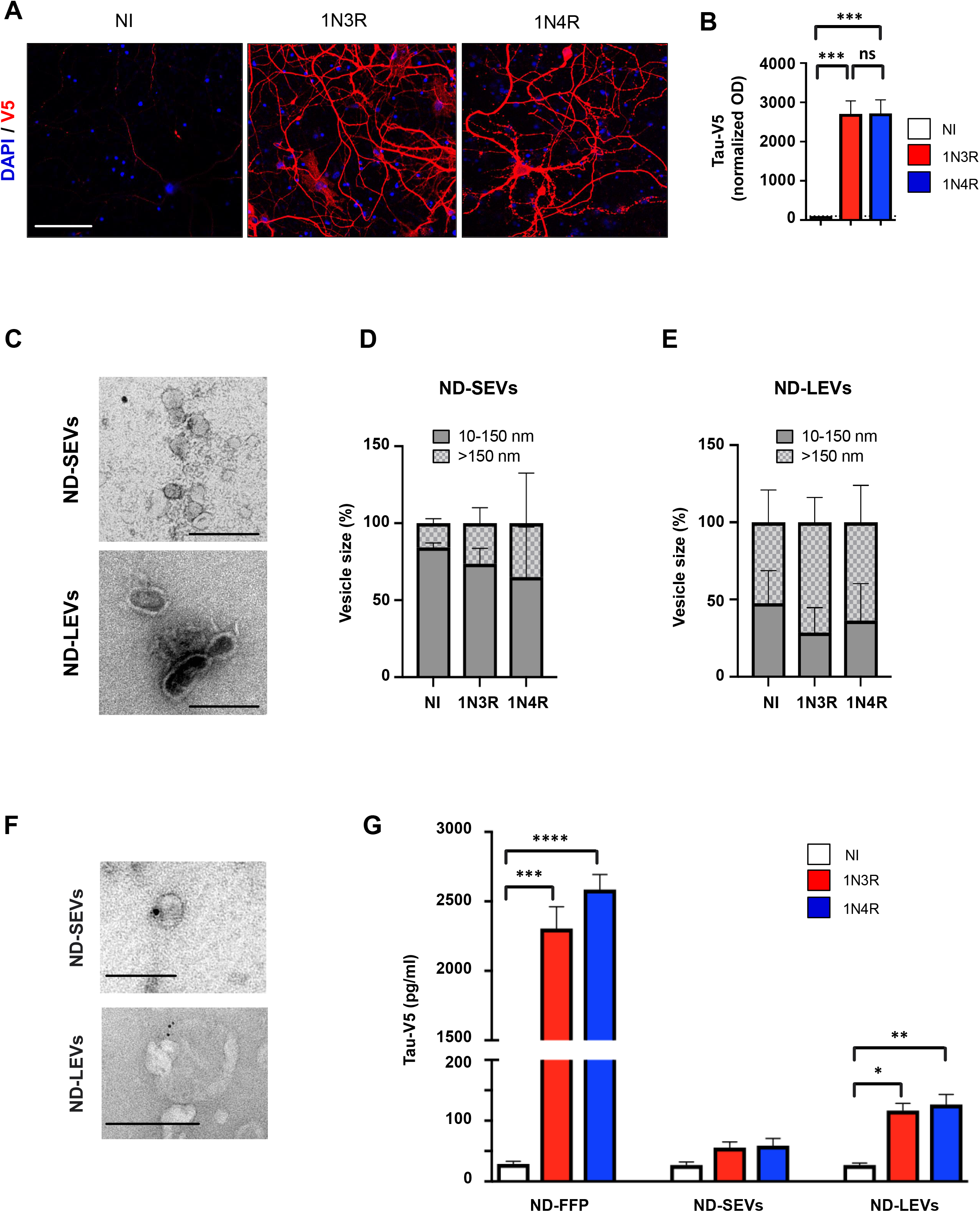
Neuronal tau is mainly secreted in a free form. **(A)** Confocal micrographs showing tau (V5+) in rat hippocampal neurons infected by LVs overexpressing 1N3R or 1N4R tau; NI=noninfected. **(B)** Histogram showing Tau-V5 accumulation (optical density normalized to that in the NI condition) in rat hippocampal neurons overexpressing 1N3R or 1N4R tau. **(C)** Electron microscopy images of ND-SEVs and ND-LEVs isolated from cultured neurons. The scale bar is 250 nm. **(D)** Histogram showing the percentage of ND-SEVs with a size between 100-150 nm and greater than 150 nm. **(E)** Histogram showing the percentage of ND-LEVs with a size between 100-150 nm and greater than 150 nm. **(F)** Electron microscopy and immunogold labeling (Cter-tau) of ND-SEVs and ND-LEVs isolated from the supernatant of neurons overexpressing 1N4R tau. The scale bar is 100 nm for ND-SEVs and 250 nm for ND-LEVs. **(G)** Histogram showing Tau-V5 concentration in the ND-FFP, ND-SEV and ND-LEV fractions from control rat hippocampal neurons (NI) and those overexpressing 1N3R or 1N4R tau. For **B** and **G**, n□=□4 cultures per condition; ordinary one-way ANOVA with Sidak’s multiple comparison test and the nonparametric KruskalLWallis test, in **B** and **G**, respectively.

### 3.3. Tau isoforms are shuttled from neurons to astrocytes mainly by EVs

The three fractions were applied to rat hippocampal astrocyte cultures, and tau transfer was evaluated at 24 h (homemade ELISA Tau-V5 and immunofluorescence) (Figure 3A). The presence of 1N3R and 1N4R tau in astrocytes after ND-LEVs treatment was detected by immunofluorescence with antibodies against the V5 epitope and GFAP (Figure 3B), and 1N3R and 1N4R tau levels in astrocytes were quantified by ELISA (Figure 3C). Although the vast majority of tau secreted by neurons was in the free form (Figure 2G), only the human 3R isoform was detected after treatment of astrocytes with ND-FFP. In contrast, among EVs, only ND-LEVs had the ability to deliver both human 1N3R and 1N4R tau to astrocytes (Figure 3C). We then asked whether tau in ND-LEVs is transferred from neurons to astrocytes more efficiently than ND-FFP. We normalized the [Tau-V5] in astrocytes to the [Tau-V5] in the fractions applied to astrocytes and secreted by neurons (ND-FFP and ND-LEVs). The percentage of tau taken up by astrocytes was significantly different between the ND-FFP and ND-LEVs groups, and tau accumulation was not significantly affected by the tau isoform (Figure 3D). Altogether, these results highlight the major role of LEVs in tau transfer between neurons and astrocytes.

**Figure 3.**
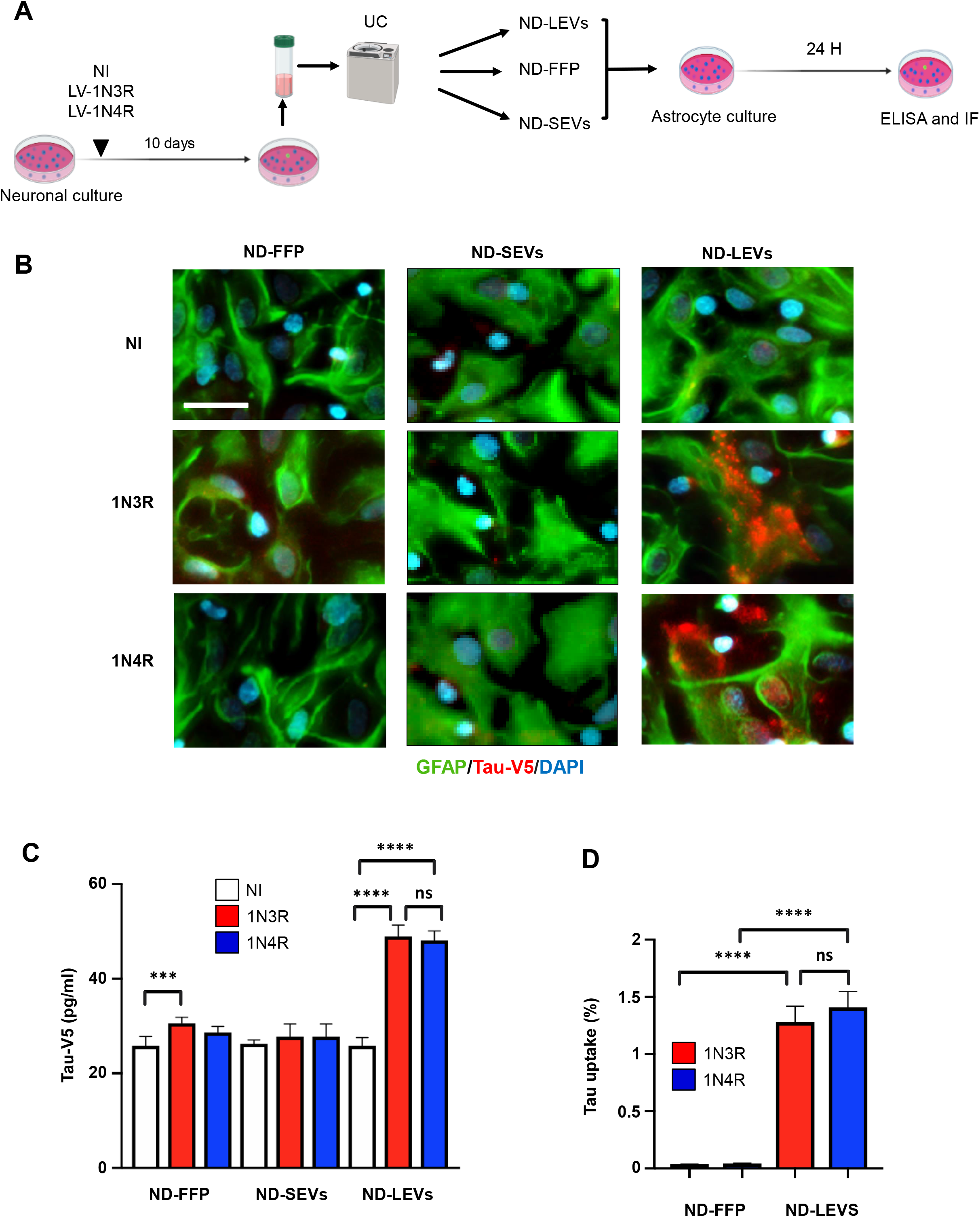
Tau is shuttled from neurons to astrocytes via LEVs. **(A)** Schematic representation of the protocol used to isolate ND-EVs from neurons infected by LVs overexpressing 1N3R or 1N4R tau. ND-EVs from NI cultures were used as controls. **(B)** Example of confocal images showing the transfer of ND-FFP, or tau in ND-SEVs and ND-LEVs from neurons overexpressing 1N3R or 1N4R tau. The scale bar is 25 µm. **(C)** Histogram showing the Tau-V5 concentration in astrocytes 24 h after incubation with the ND-FFP, ND-SEVs and ND-LEVs fractions from control rat hippocampal neurons (NI) and those overexpressing 1N3R or 1N4R tau. **(D)** Histogram showing the tau uptake efficiency 24 h after incubation with the ND-FFP and ND-LEVs fractions from control rat hippocampal neurons (NI) and those overexpressing 1N3R or 1N4R tau. For **C** and **D**, n□=□4 cultures per condition; ordinary one-way ANOVA with Sidak’s multiple comparison test.

### 3.4. 3R and 4R tau-accumulating LEVs treatments induce differential mitochondrial consequences in astrocytes

We previously demonstrated that the accumulation of different human tau isoforms in astrocytes differentially affects the astrocytic mitochondrial system [24]. The effect of ND-LEVs derived from neurons accumulating 3R or 4R tau was assessed by following a Mitotimer biosensor with high-content live microscopy (Figure 4A). This multiparametric approach can reveal changes in individual mitochondria over several hours/days [36]. Mitochondrial features (including the redox state, morphology, and dynamic changes) were measured before (BL) ND-LEVs^CFP^, ND-LEVs^3R^ or ND-LEVs^4R^ treatment and at 6 h and 24 h after treatment. For each astrocyte- and each mitochondria-related criterion, the change score was calculated relative to the BL value and normalized to that of the control group (ND-LEVs^CFP^).

**Figure 4.**
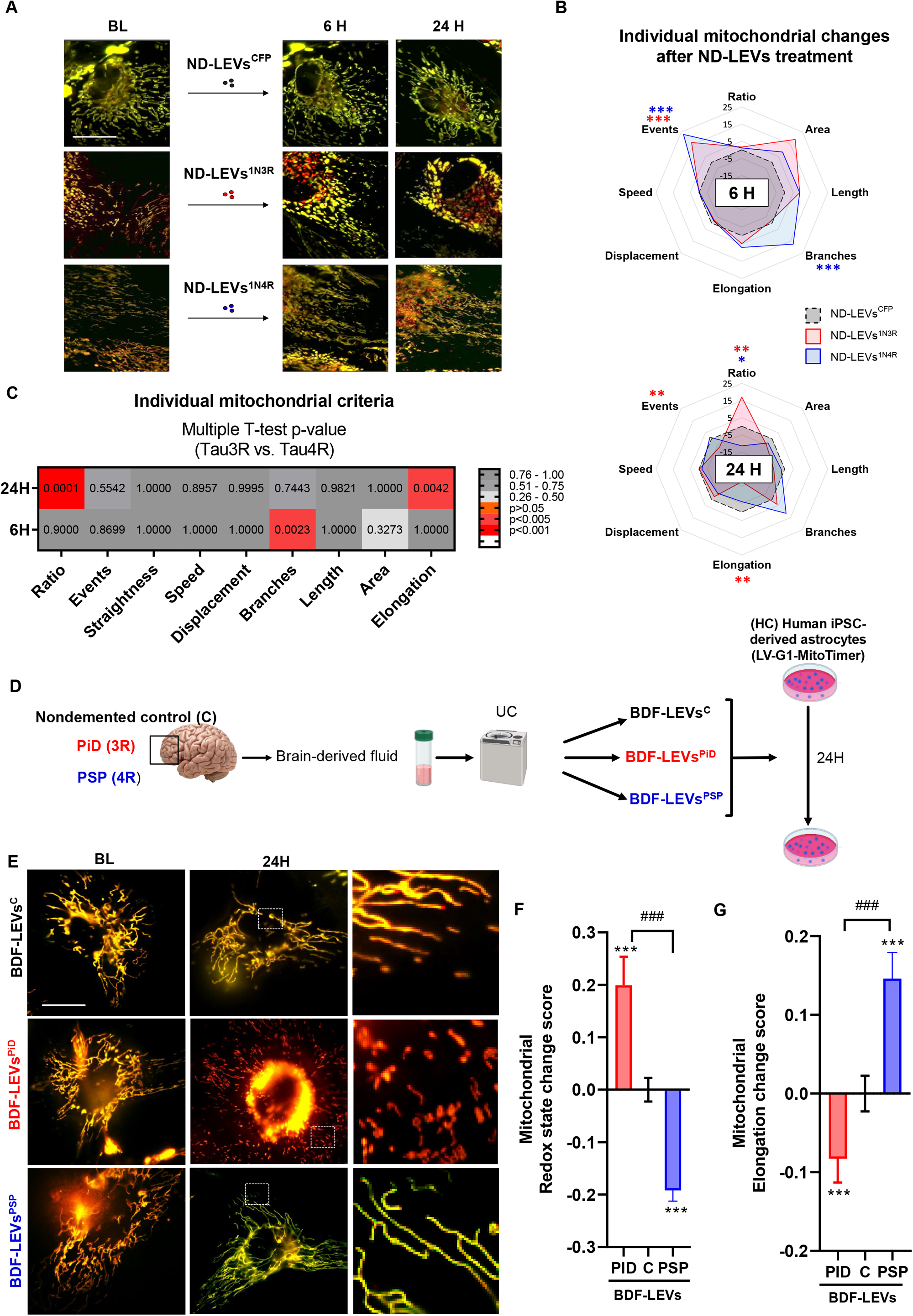
Tau isoform-containing LEVs induce mitochondrial dysfunction in astrocytes. **(A)** Micrographs of the mitochondria labeled with the biosensor mitotimer before (BL) and after (6 h and 24 h) treatment with ND-LEVs^CFP^, ND-LEVs^1N3R^ and ND-LEVs^1N4R^. **(B)** Radar charts showing mitochondrial morphology, redox state and mobility changes (normalized to BL values and the value of the ND-LEV^CFP^ group) in astrocytes treated with ND-LEVs^1N3R^ and ND-LEVs^1N4R^. Two-way matched ANOVA followed by post hoc analyses (Tukey’s test) was used to compare the effect of each treatment to that of ND-LEVs^CFP^ (_∗_*p* < 0.05, _∗∗_*p* < 0.01 and _∗∗∗_*p* < 0.001). **(C)** Heatmap showing the p value determined by multiple t tests assessing the difference in mitochondrial changes 6 h and 24 h after treatment with ND-LEVs^1N3R^ and ND-LEVs^1N4R^. **(D)** Schematic representation of the protocols used to isolate BDF-EVs from patients diagnosed with PiD or PSP and non-demented control subjects (C) and to treat iPSC-derived astrocytes. **(E)** Micrographs of the mitochondrial system in human iPSC-derived astrocytes expressing the biosensor mitotimer before (BL) and after (6 h and 24 h) treatment with BDF-LEVs^PiD^ and BDF-LEVs^PSP^. **(F)** Histogram showing mitochondrial redox state changes in iPSC-derived astrocytes 24 h after BDF-LEVs treatment. **(G)** Histogram showing mitochondrial elongation in iPSC-derived astrocytes 24 h after BDF-LEVs treatment. N□=□cultures/cells/mitochondria; ND-LEVs^CFP^: 3/21/801, ND-LEVs^1N3R^: 3/15/958, ND-LEVs^1N4R^: 4/14/789, BFD-LEVs^C^: 3/19/3883, BFD-LEVs^PiD^: 3/21/749, BFD-LEVs^PSP^: 4/17/2555. Two-way matched ANOVA followed by post hoc analyses (Tukey’s test) were used to compare the effect of treatments with that of the other treatments (###p < 0.001) or with the nondemented control group (_***_*p* < 0.001). Scale bars are 30□µm.

Six hours after ND-LEVs^3R^ or ND-LEVs^4R^ treatments, we observed that the number of mitochondrial events (fusion/fission) significantly increased compared to the control condition ND-LEVs^CFP^ (Figure 4B). However, only ND-LEVs^4R^ increased the morphological complexity by significantly increasing the number of branches of mitochondria (Figure 4B). At 6H, multiple t-tests between treatments indicated that the mitochondrial branch criteria distinguish between the two types of ND-LEVs^3R^ or ND-LEVs^4R^ treatments (Figure 4C). Twenty-four hours after ND-LEVs^3R^ or ND-LEVs^4R^ treatments, we observed that ND-LEVs^3R^ significantly increases the redox state of mitochondria and reduces the number of events and the elongation of astrocyte mitochondria (Figure 4B). Conversely, ND-LEVs^4R^ reduces the redox state and tends to more branched mitochondria. Multiple t-tests between treatments revealed that the mitochondrial redox state and elongation significantly differ between ND-LEVs^3R^ or ND-LEVs^4R^ treatments (Figure 4C). These data suggest that the mitochondrial system of astrocytes is rapidly sensitive (6h) to the pathological content of LEVs and may be damaging in the longer term (24H).

Then, we evaluated changes in mitochondrial parameters (24 h) in human iPSC-derived astrocytes (expressing the biosensor Mitotimer) treated with BDF-LEVs isolated from a pool of patients diagnosed with 3R-tau (PiD), 4R-tau (PSP) tauopathy or non-demented control subjects (C) (Figure 4E, Figure S2A). The size, concentration, and protein content of BDF-LEVs were initially measured using NTA and chromatography-tandem mass spectrometry (LCLMS/MS) (Figure S2A-C). Regarding redox state, astrocytes treated with BDF-LEVs^PID^ exhibited significantly increased mitochondrial oxidation, whereas mitochondrial oxidation was significantly reduced in astrocytes treated with BDF-LEVs^PSP^ (Figure 4E, F). Moreover, BDF-LEVs^PID^ significantly reduced mitochondrial elongation, whereas the mitochondria of astrocytes treated with BDF-LEVs^PSP^ appeared significantly more elongated and widely distributed in the cell (Figure 4E, G). Altogether, these results revealed that EVs from cells with 3R tau accumulation had deleterious effects on mitochondrial redox state and morphology, while EVs from cells with 4R tau accumulation induced filamentation and had less severe functional effects on the astrocytic mitochondrial system.

## 4. Discussion

This study investigates the involvement of glial cells and EVs in tau spreading. To date, most studies addressing the spreading process have focused on neurons. These cells are indeed the most affected cells in tauopathies, especially in AD, the main feature of which is the aggregation of tau protein (3R and 4R tau) into paired helical filaments within neurons. Nevertheless, tau inclusions are also found in glial cells, including astrocytes, in other primary tauopathies. Astrocytes are the principal glial cells in the brain and play a supporting role in supplying energy and nutrition to neurons. Evidence shows that astroglial atrophy occurs in early stages of neurodegeneration, potentially leading to disruption in synaptic connectivity [38]. Astrocytes form a direct link between pre- and postsynaptic neurons and may therefore be involved in the propagation of tau between interconnected brain regions. Moreover, tau in astrocytes involved in tripartite synapses can impair synaptic function, as has been described in the brain of AD patients [39].

Here, we evaluated whether tau protein is shuttled from neurons to astrocytes via EVs to provide a potential explanation for the accumulation of tau in astrocytes in the AD brain [24]. We decided to first investigate the related cellular mechanisms in primary neurons overexpressing either 3R or 4R tau. EVs from murine primary neurons expressing human 3R or 4R tau were isolated and applied to murine primary astrocytes. Whereas tau is mainly secreted in a free form, EVs equally transfer both the two isoforms to astrocytes. In agreement with the work of Mate de Gerando and colleagues [15], we confirmed that tau is transferred from neurons to astrocytes. We also validated the dependence of astrocytes on neurons for tau spreading [40]. Most importantly, through a direct comparison of free and EVs-associated tau, we validated the high potential of EVs to shuttle tau between neurons and astrocytes. This comparison also highlights the importance of brain-derived EVs in spreading tau pathology in Alzheimer’s disease [21,22].

We then studied the effects of EV-mediated 3R or 4R tau transfer on astrocytes. Our data demonstrated that the transfer of EVs from neurons with 3R or 4R tau accumulation induced precise changes in the astrocytic mitochondrial system. Indeed, EVs originating from neurons with 3R tau accumulation rapidly show features of mitochondrial dysfunction/damage (a transient increase in the number of events that result in fragmentation and a substantial rise in the mitochondrial redox state). In contrast, astrocytes treated with EVs originating from neurons with 4R tau accumulation show features of transient adaptation/compensation of the mitochondrial system (transient ramification, a reduction in the redox state and/or an increase in mitochondrial turnover) [41,42]. Similar results were obtained after overexpression of 3R and 4R tau in astrocytes [24], whereas opposite effects were observed in neurons [43].

Numerous studies have already shown that the accumulation of abnormal tau protein impairs mitochondrial function, which leads to cell degeneration. These alterations in function include mitochondrial transport, dynamics, bioenergetics and mitophagy [44].

Initially, because tau is a microtubule-binding protein, mitochondrial disturbances due to abnormal tau have long been attributed to a change in the organization of the cytoskeletal network [45]. However, recently, several studies demonstrated that abnormal tau has a direct effect on mitochondria via binding. Indeed, it has been shown that abnormal tau can alter MFN2 levels and trigger the mislocalization and clustering of DRP1, triggering the elongation of mitochondria [46]. Abnormal tau can also directly interfere with the Parkin protein, inhibit mitophagy [47], and significantly reduce the activity of complex I, voltage-dependent anion channel (VDAC) and respiratory complex V subunits [48]. Abnormal tau protein may also impact ER-mitochondrial coupling, which could affect all mitochondrial functions [49]. The vast majority of related studies have used neurons and 4R tau isoforms. Consequently, the importance of the specific tau isoform in determining the extent of mitochondrial alterations is still unclear.

Our studies demonstrated that 3R and 4R tau are transferred from neurons to astrocytes with the same efficiency. We can hypothesize that 3R and 4R tau act on astrocytic mitochondria through different molecular mechanisms. First, whether three or four microtubule-binding domains are present slightly influences the binding affinity of tau to microtubules. Indeed, 4R tau binds to microtubules with a 3-fold higher affinity than tau [50]. We can also assume, as demonstrated in neurons, that tau protein may bind differently to filamentous actin to induce the formation of aligned bundles of actin filaments, therefore modifying the organization of the cytoskeletal network [45]. Second, because it is more soluble and has stronger kinesin inhibitory activity than 4R tau, 3R tau may induce strong steric inhibition of the binding of mitochondria to microtubules, leading to their immobilization [51]. Furthermore, the presence of one cysteine (C322) in 3R tau allows the formation of intermolecular bridges, whereas the presence of two cysteines (C291, C322) in 4R tau leads mainly to the formation of intramolecular bridges [52]. These differences in the ability to form inter- or intradisulfide bridges may explain why 3R tau more readily aggregates to form oligomers and polymers than 4R tau. If these differences in properties are maintained in EVs, they could explain the differential effects of 3R and 4R tau on mitochondria observed in our study. Indeed, EVs carrying 3R tau may contain more tau peptides that are directly toxic to mitochondria [21,22,44] than EVs containing 4R tau. To extrapolate this effect to the human condition, we next isolated EVs from the brain fluids of patients with a 3R tau-related primary tauopathy (PiD) and a 4R tau-related primary tauopathy (PSP). Recently, we and others showed that EVs isolated from tauopathy patient-derived brain fluids contain tau seeds that might be involved in tau spreading [21]. Application of these EVs to iPSC-derived astrocytes validated their deleterious effect on mitochondria and showed that EVs from pure 3R tau-related tauopathy samples are the most aggressive, confirming our data from murine primary cultures. Altogether, our data are consistent with our previous study showing that 3R and 4R tau overexpression in astrocytes differentially alters the mitochondrial localization, trafficking and function of these tau isoforms, suggesting that astrocytes may play a more substantial role than expected in AD [24] and other pure tauopathies. However, it is essential to mention that the observed effects on mitochondria may be more complex than the effect of tau alone. Indeed, EVs carry many other proteins (or protein peptides) and mRNAs that could impact the astrocytic mitochondrial system. Recent evidence suggests that the transfer of mitochondrial content by EVs modifies metabolic and inflammatory responses in recipient cells [53]. Alone or in combination, these effects are consistent with our observations of altered mitochondrial transport and dynamics after EVs-mediated 3R tau transfer.

Here, we found that human-derived EVs play a role in astrocytic dysregulation. We also described a new potential pathway for tau spreading between interconnected regions surrounding the synaptic cleft. We showed that EVs play a significant role in propagating 3R and 4R tau to astrocytes. Treatment with accumulated tau-containing EVs originating from neurons with 3R tau accumulation disturbed the astrocytic mitochondrial system and had very damaging effects. This study raises many questions about strategies for clearing free extracellular tau. However, targeting pathological EVs is very challenging, and further studies are required to characterize EVs cargos/transport, as recently described [54-56]. This will help not only in differentiating tauopathies but also in designing specific tools to block tau spreading. To conclude, we highlight new mechanisms that explain how pathology can spread from neurons to surrounding astrocytes and alter their functioning.

## Supplementary materials

**Figure S1. BFD-LEVs cell toxicity assay**. Histogram showing the number of astrocytes still present in the well (counting of DAPI+ cells) 24 h after treatment with BDF-LEVs isolated from non-demented control subjects (C), PiD patients and PSP patients. BDF-LEVs were administered at doses of 10^5^, 10^6^ and 10^7^ BDF-EVs/astrocyte. NL=L5-7 cultures per condition; ordinary one-way ANOVA with Sidak’s multiple comparison test.

**Figure S2. BFD-LEVs characterization and quality control. (A)** Table showing the number of vesicles per ml, the level of hTau per vesicle and the level of hTau per vesicle normalized to tissue weight for each pooled sample (C, PiD and PSP). **(B)** Histogram of the size distribution of BFD-LEVs (between 10 and 150 nm or larger than 150 nm). **(C)** Heatmap depicting the relative abundances of proteins in different GOCC categories characterized as noncontaminant or potential contaminant in BFD-LEVs. **(D)** Histogram depicting the relative abundances of proteins in different MISEV18 categories characterized as EVs-associated or non-EVs-associated.

## Acknowledgments and funding sources

This research was funded by grants from the Investissement d’Avenir LabEx (Investing in the Future Laboratory Excellence) program, DISTALZ (Development of Innovative Strategies for a Transdisciplinary Approach to ALZheimer’s disease), Fondation France Alzheimer (project: EV-Tau and spreading), Fondation Alzheimer (project Ectausome), Fondation pour la Recherche Médicale, ANR grants (GRAND, TONIC, TAUSEED), and the PSP France Association. Our laboratories were also supported by LiCEND (Lille Centre of Excellence in Neurodegenerative Disorders), CNRS, Inserm, Métropole Européenne de Lille, the University of Lille, I-SITE ULNE, Région Hauts de France and FEDER. We are grateful to the Lille Neurobank and Pr Claude-Alain Maurage for access to the human brain samples. This study was also supported by a Synapsis Foundation and the Lausanne University Hospital (CHUV). The authors thank the Protein Analysis Facility of the University of Lausanne, particularly Dr. M. Quadroni, for technical support. We are grateful to the UMS-2014 US41 PLBS for access to the confocal microscopy at the HU site of the BioImaging Center Lille and for their help.

## Author contributions

Designed and conceptualized the study: K.R., M.C. and L.B. Writing, review and editing: K.R., M.C. and L.B. Support for manuscript editing: R.P., E.L., N.D. and N.T. Experimentation: R.P., J.E., E.L., J.H.L., V.Z., L.C., S.P., R.D.P., S.B., and V.D. Funding acquisition: K.R., M.C. and L.B. Supervision: K.R. and M.C. All authors read and approved the final manuscript.

## Ethics approval and consent to participate

The study was conducted in accordance with the Declaration of Helsinki. Brain extracts were obtained from the Lille Neurobank (fulfilling French legal requirements concerning biological resources and declared to the competent authority under the number DC-2008-642) with donor consent and ethics committee approval in compliance with data protection regulations. For human- or murine-derived primary cultures, experiments were performed according to an ethics protocol approved by our institutional review committee (CER-VD 2018-01622, Lausanne, Switzerland and APAFIS#2264-2015101320441671 from CEEA75, Lille, France).

## Conflict of interest

The authors declare no competing interest related to funding.

## Notes

### Competing Interest Statement

The authors have declared no competing interest.

