## Supplementary figures and images for "Accumulation of Tau in Extracellular Vesicles Disturbs the Astrocytic Mitochondrial System"

### Figure S1

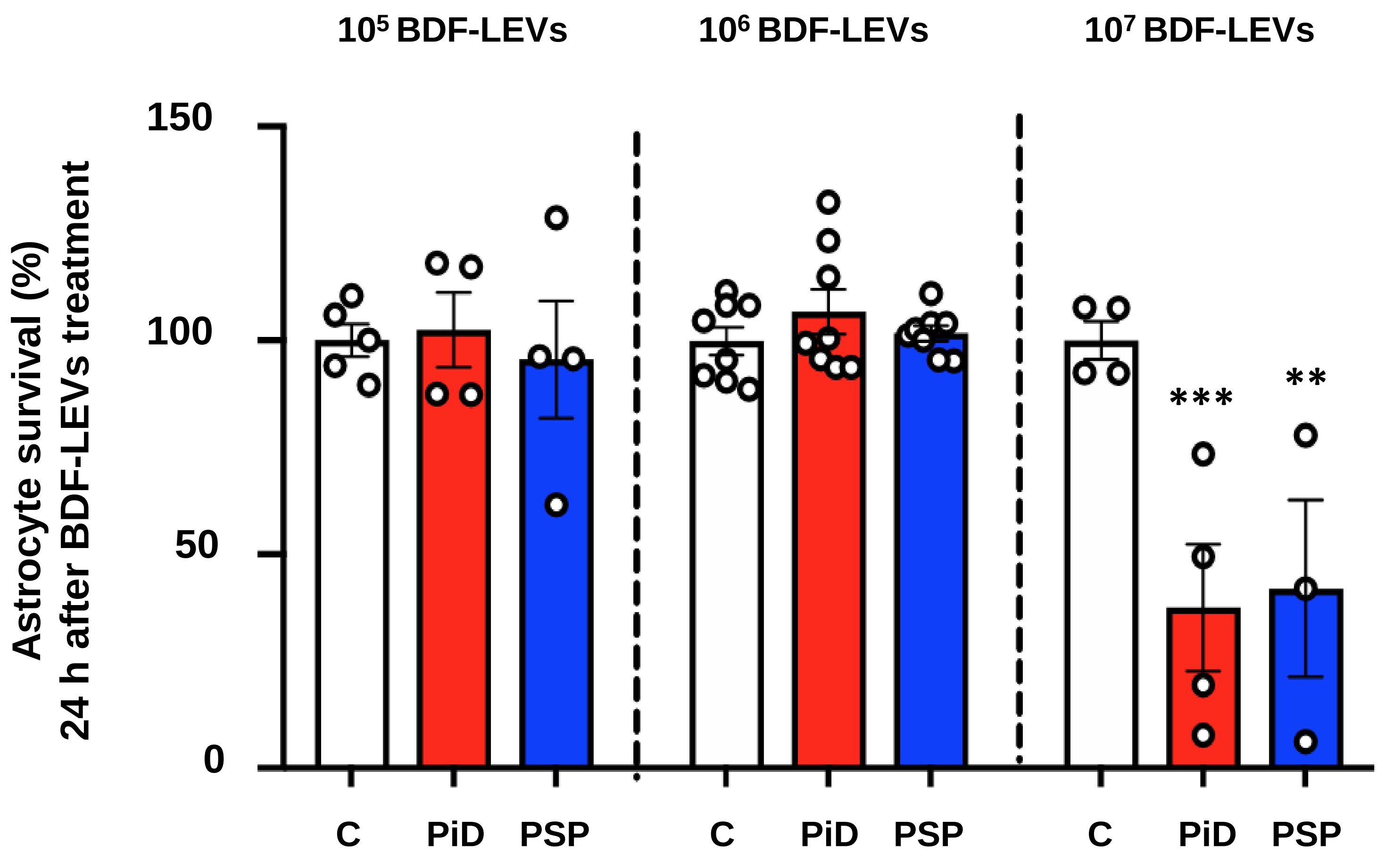

### Figure S2

A

| | C | PiD | PSP |
|---|---|---|---|
| **Vesicles/ml** | $1.04x10^{13}$ | $2.03x10^{12}$ | $9.17x10^{12}$ |
| **hTau/vesicle** | $3.71x10^{-8}$ | $4.2x10^{-8}$ | $3.6x10^{-8}$ |
| **hTau/vesicle/g tissue** | $4.73x10^{-9}$ | $5.35x10^{-9}$ | $3.62x10^{-9}$ |

B

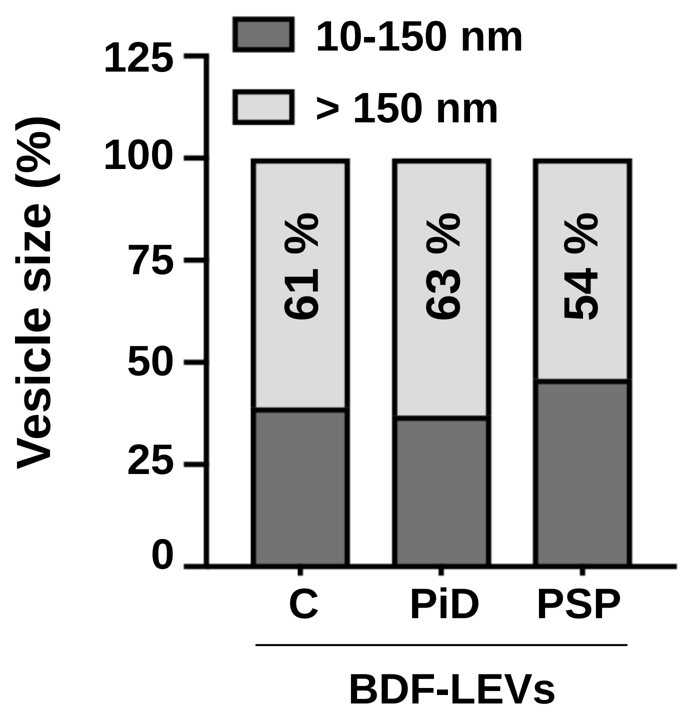


C

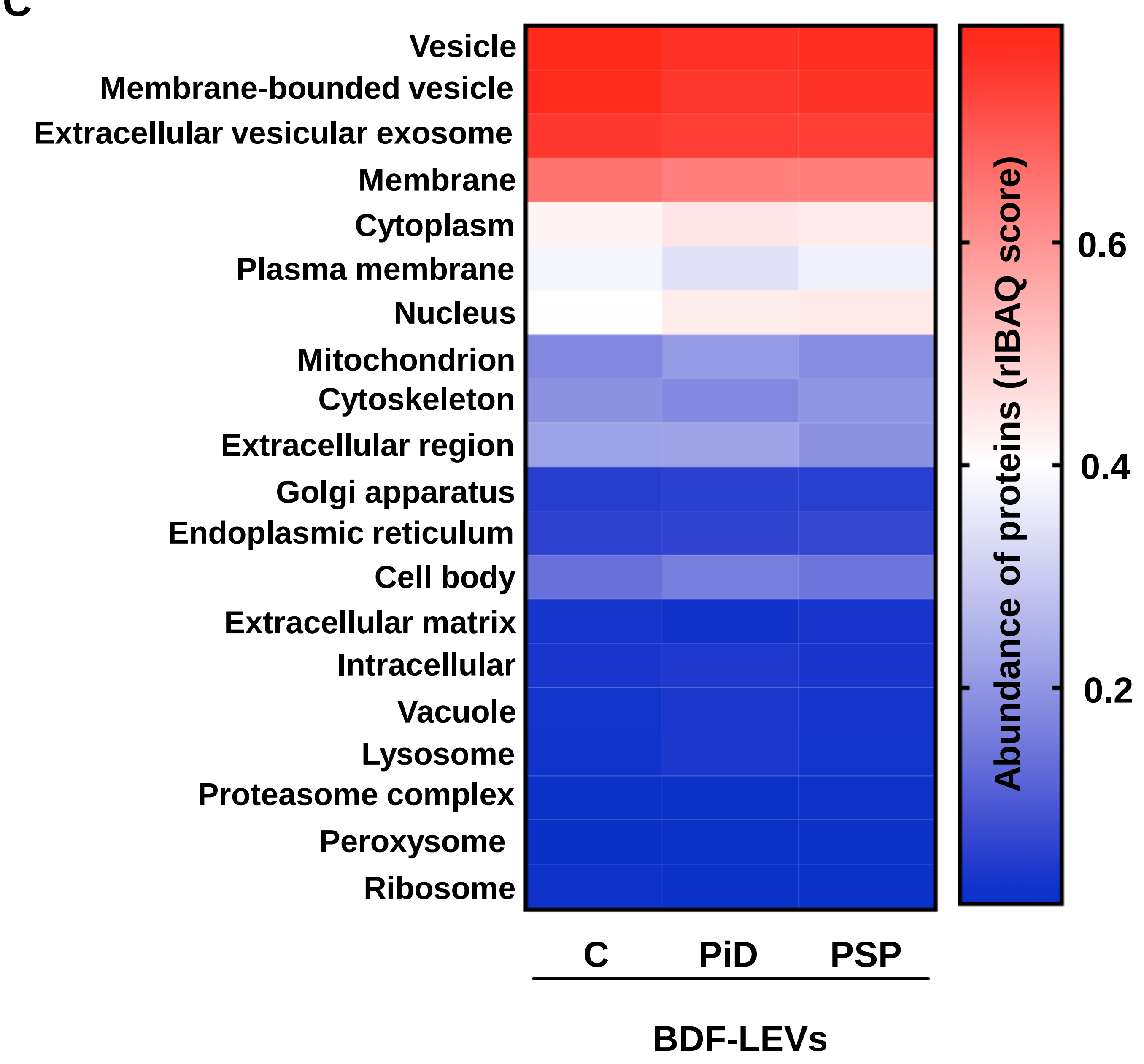


D

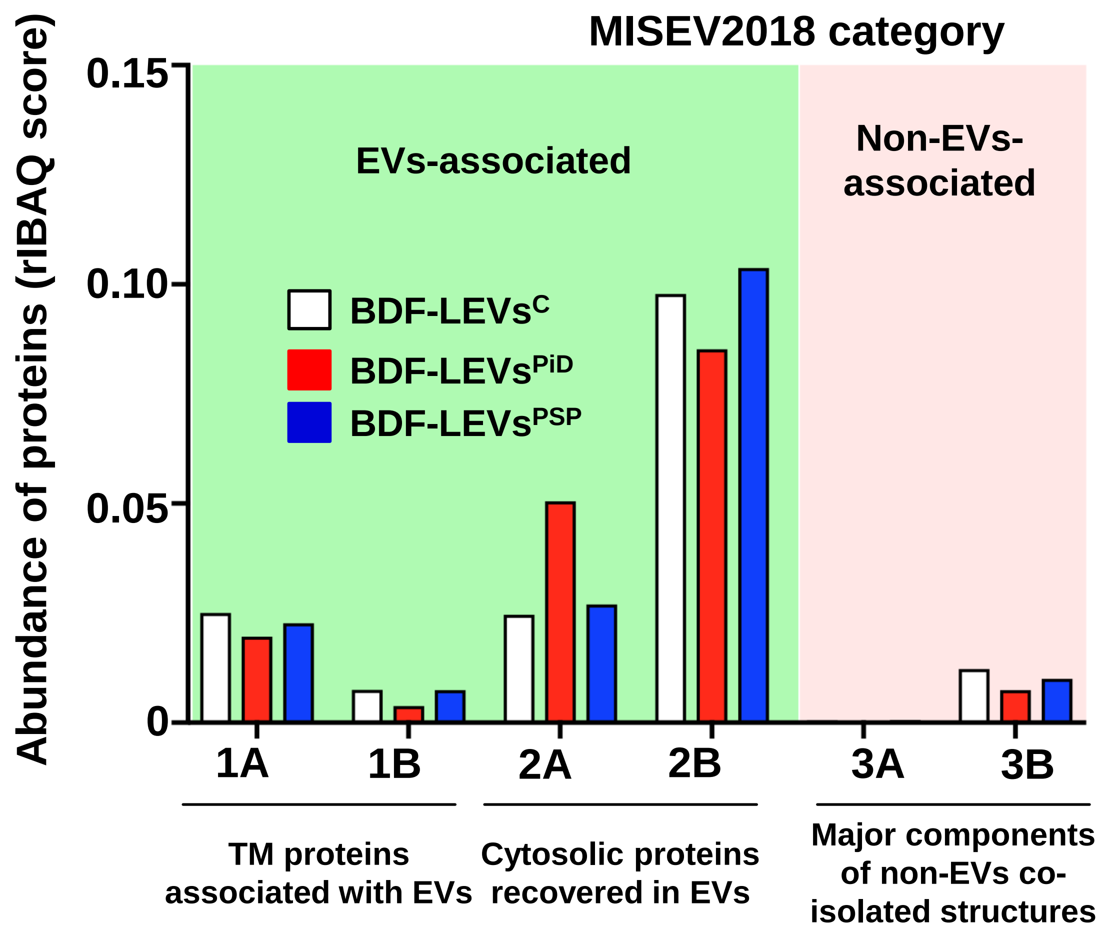
